# Odor Modulates the Temporal Dynamics of Fear Memory Consolidation

**DOI:** 10.1101/2019.12.19.881615

**Authors:** Stephanie L. Grella, Amanda H. Fortin, Olivia McKissick, Heloise Leblanc, Steve Ramirez

## Abstract

Systems consolidation (SC) theory proposes that recent, contextually rich memories are stored in the hippocampus (HPC). As these memories become remote, they are believed to rely more heavily on cortical structures within the prefrontal cortex (PFC), where they lose much of their contextual detail and become schematized. Odor is a particularly evocative cue for intense remote memory recall and despite these memories being remote, they are highly contextual. In instances such as post-traumatic stress disorder (PTSD), intense remote memory recall can occur years after trauma, which seemingly contradicts SC. We hypothesized that odor may shift the organization of salient or fearful memories such that when paired with an odor at the time of encoding, they are delayed in the de-contextualization process that occurs across time, and retrieval may still rely on the HPC, where memories are imbued with contextually rich information, even at remote time points. We investigated this by tagging odor- and non-odor-associated fear memories in male c57BL/6 mice and assessed recall and *c-Fos* expression in the dorsal CA1 (dCA1) and prelimbic cortex (PL) 1 d or 21 d later. In support of SC, our data showed that recent memories were more dCA1-dependent whereas remote memories were more PL-dependent. However, we also found that odor influenced this temporal dynamic biasing the memory system from the PL to the dCA1 when odor cues were present. Behaviorally, inhibiting the dCA1 with activity-dependent DREADDs had no effect on recall at 1 d and unexpectedly caused an increase in freezing at 21 d. Together, these findings demonstrate that odor can shift the organization of fear memories at the systems level.

## INTRODUCTION

Remembering personal experiences, or episodic memories, relies upon the integrity of the hippocampus (HPC) (Scoville and Milner, 1957). When an episodic memory is formed, many of the contextual elements of the experience are encoded. The set of brain cells, or neuronal ensembles active during memory formation can be referred to as an engram (Semon, 1921) and natural recall of an episodic memory involves reactivation of those engrams (Deng et al., 2013; Tonegawa et al., 2015; Holtmaat and Caroni, 2016). Experiential recall can also be induced by artificial reactivation of engram cells using genetic tagging and optogenetic or chemogenetic stimulation.

New memories which, are initially labile, gain stability and permanence through consolidation. Consolidation is a process of reorganization that occurs within the hours following the encoding of an experience at the synaptic level (local changes in connectivity), and gradually over years at the systems level (brain-wide changes in connectivity) (Frankland and Bontempi, 2005). Some of the first evidence that led researchers to understand the fundamental necessity of the HPC in the early stages of memory formation and consolidation came from individuals with hippocampal damage who experienced anterograde amnesia or the inability to form new memories, as a result. These patients also showed temporally graded retrograde amnesia (sometimes called the Ribot gradient) (Ribot, 1882) where the magnitude of hippocampal damage was correlated with the temporal gradient of amnesia. In these individuals, damage to the HPC was associated with deficits for recently acquired memories, however, more distant memories were still intact (Zola-Morgan and Squire, 1986; Rempel-Clower et al., 1996; Kapur and Brooks, 1999; Bayley et al., 2003; Kirwan et al., 2008). The opposite pattern was observed when cortical structures like the PFC sustained damage - remote memories were lost (retrograde amnesia), while recent memory was unaffected (Markowitsch et al., 1993; Mangels et al., 1996; Reed and Squire, 1998; Murre et al., 2001; Bayley et al., 2003, 2005; Squire and Bayley, 2007; Squire and Wixted, 2011).

The theory of systems consolidation (SC) proposes a mechanism for these findings, arguing that as memories becomes older, or more remote, they become less HPC-dependent, and more dependent on the PFC (Ribot, 1882; Squire, 1992; Lechner et al., 1999; Frankland et al., 2004; Wiltgen et al., 2004; Frankland and Bontempi, 2005; Nadel et al., 2007; Squire and Bayley, 2007; Squire et al., 2015), a process that is thought to be mediated by post-learning spontaneous engram reactivation during sleep (Hebb, 1949; Marr, 1971; Skaggs and McNaughton, 1996; Redish and Touretzky, 1998; Mölle et al., 2006; de Sousa et al., 2019). As this transition occurs, it is hypothesized that memories lose many of their rich contextual details (Tse et al., 2007; Kitamura et al., 2017). In accordance with this, retrieval of a memory shortly after an experience involves reactivation of HPC engram cells whereas retrieval occurring later involves reactivation of PFC engram cells (Maviel et al., 2004; Wiltgen et al., 2004, 2010; Ross and Eichenbaum, 2006; Gonzalez et al., 2013; Doron and Goshen, 2018). However, several observations challenge this model, such as the qualitative observation that people often retrieve remote memories that are vivid and highly detailed (Bonnici et al., 2012). Likewise, several studies have shown that there is activity in both structures during both recent and remote memory recall (Piolino et al., 2004; Goshen et al., 2011; Bonnici et al., 2012; Bonaccorsi et al., 2013; Lux et al., 2016) and that damage to the HPC sometimes affects remote memory as well as recent memory (Shimizu et al., 2000; Debiec et al., 2002; Broadbent et al., 2006; Teixeira et al., 2006; Kirwan et al., 2008; Sutherland et al., 2008; Irish et al., 2010; Zelikowsky et al., 2012).

Multiple trace theory (MTT), an alternative to the standard model of SC, accounts for these findings arguing that irrespective of when they are acquired, vivid autobiographical memories always engage the HPC (Nadel and Moscovitch, 1997; Nadel et al., 2000; Rosenbaum et al., 2001; Gilboa et al., 2004; Meeter and Murre, 2004; Moscovitch et al., 2005; Rekkas and Constable, 2005). Here, remote memories retain rich contextual detail (Suzuki and Naya, 2011) as long as the trace in the HPC is dominant at the time of retrieval. However, there is evidence to suggest that engrams are in fact, encoded in parallel across both brain regions (Kirchhoff et al., 2000; Blumenfeld and Ranganath, 2007; Qin et al., 2007; Goshen et al., 2011; Lesburgueres et al., 2011; Tayler et al., 2013) and that the architecture of memory traces formed during encoding are sparse but distributed (Rao-Ruiz et al., 2019; Roy et al., 2019) supporting a third, more flexible model, competitive trace theory (CTT). CTT suggests that multiple traces exist (e.g., HPC and PFC) and that memories do become more schematized or decontextualized over time and more reliant on neocortical storage but that the HPC can function to recontextualize memories, at any time, even at remote time points (Yassa and Reagh, 2013).

The temporal separation between recent and remote memories is ambiguous (Doron and Goshen, 2018). In the case of post-traumatic stress disorder (PTSD) these lines may be even more blurred given that sometimes trauma-related memories can be recalled with vivid detail many years after the traumatic event (Wagenaar and Groeneweg, 1990; Schelach and Nachson, 2001; Vermetten and Bremner, 2003; Rubin et al., 2004, 2011; Berntsen and Thomsen, 2005; Janssen et al., 2011; Fernández-Lansac and Crespo, 2017) which may be related to the highly arousing nature of these memories (Kensinger and Schacter, 2008; Liu and McNally, 2017), however, there are conflicting views regarding how emotion modulates memory in general (Rubin et al., 2008; Fernández-Lansac and Crespo, 2017). More specifically, odor seems to be a particularly powerful contextual cue that can evoke this experience long after memories are stabilized (Vermetten and Bremner, 2003; Arshamian et al., 2013). Odors can be potent triggers of memories, (Daniels and Vermetten, 2016) and as such, we believe that if they are present during encoding, they can shift the organization of arousing memories at the systems level, to a state of HPC-dependent processing.

To analyze the modulatory role of odor in the temporal dynamics of memory consolidation, we designed an experiment to visualize sets of engram cells at recent and remote time points. We hypothesized that fear related memories would become more PFC dependent and less HPC dependent with the passage of time. Secondly, we hypothesized that if we fear-conditioned animals with an odor present, that this contextual element would shift the temporal organization of that memory such that it would be less reliant on the PFC, and more so on the HPC even at remote time points. Theoretically, if our data supports SC then inhibiting activity in the HPC should only affect recent and not remote memory retrieval. However, if odor can shift this organization then inactivation of HPC engram cells related to an odor-associated fear memory will also affect retrieval at the remote time point. If our data support MTT or CTT, then we should also see an impairment also at both time points.

Briefly, we utilized chemogenic and virus-based strategies to locate and tag cells during the formation of a contextual fear memory in c57BL/6 wildtype mice in a doxycycline (DOX)-regulated manner. The viruses used comprise an inducible and activity-dependent system in order to control which cells are tagged and later inhibited. It is inducible because it is controlled by DOX (a derivative of tetracycline), which when present in the animal’s diet prevents expression of the inhibitory hM4Di DREADD and the associated fluorescent reporter eYFP, and when absent allows for their expression. The system is activity-dependent because the expression of these proteins is driven by the immediate early gene *c-Fos* promoter, often used as a neuronal marker of activity. Mice were taken off-DOX and then fear-conditioned to tag the cells involved. Mice were placed back on DOX and then tested for memory-recall either 1 d or 21 d later and half the mice were exposed to almond extract odor during conditioning and recall. Upon recall, mice were given CNO to inhibit the cells active during fear conditioning. We measured freezing levels across sessions and at the end of the experiment we quantified *c-Fos* activity in the dorsal CA1 (dCA1) of the HPC as well as the prelimbic cortex (PL) of the PFC.

## RESULTS

### Viral Transduction and Activity-Dependent Engram Labeling

Viral transduction was localized to target regions (PL and dCA1) (Figure 1A-C). Mis-targeted animals were excluded from data analysis. There were no group differences in the proportion of DAPI-labeled cells within the PL (Figure 1E&G left) and dCA1 (Figure 1D&F left). Likewise, there were no group differences in the percentage of cells that were labeled with eYFP during fear conditioning in the PL (Figure 1E&G right) and dCA1 (Figure 1D&F right).

**Figure 1.**
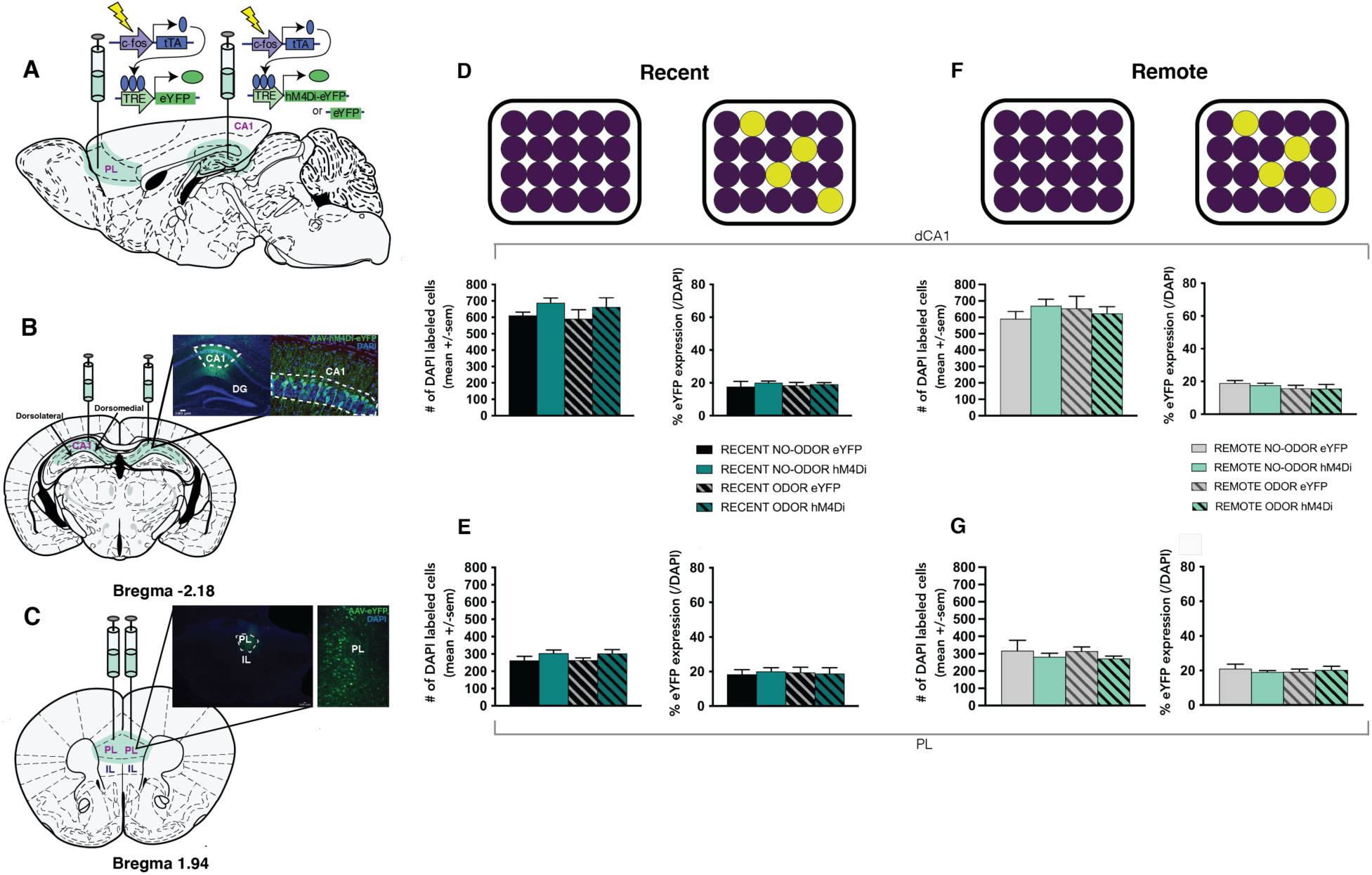
Viral Transduction and Activity-Dependent Engram Labeling. **(A-C)** Viral transduction was localized to target regions: Prelimbic Cortex (PL) and dorsal CA1 (dCA1). In the PL, all mice were injected with AAV9-cFos-tTa-TRE-eYFP and in the dCA1 with either AAV9-cFos-tTa-TRE-hM4Di-eYFP (inhibitory DREADDs) or AAV9-cFos-tTa-TRE-eYFP. The proportion of DAPI-labeled cells **(left)** and eYFP+ cells **(right)** labeled during fear conditioning, when fear memory recall was tested after 1 day (Recent condition), within the dCA1 **(D)** and the PL **(E)**. The proportion of DAPI-labeled cells **(left)** and eYFP+ cells **(right)** labeled during fear conditioning, when fear memory recall was tested after 21 days (Remote condition), within the dCA1 **(F)** and the PL **(G)**. Groups: RECENT-NO-ODOR-eYFP (n=7), RECENT-NO-ODOR-hM4Di (n=5), RECENT-ODOR-eYFP (n=7), RECENT-ODOR-hM4Di (n=5), REMOTE-NO-ODOR-eYFP (n=8), REMOTE-NO-ODOR-hM4Di (n=5), REMOTE-ODOR-eYFP (n=7), REMOTE-ODOR-hM4Di (n=5). p< 0.05, ** = p<0.01, *** = p<0.001.

### Inhibiting the CA1 with DREADDs did not Impair Fear Memory Recall

To assess whether odor modulates systems consolidation of a fear memory we employed a four-shock contextual fear conditioning protocol. Animals were initially placed in the context without shock for 198s. Figure 2B shows that during this PRE-Shock period the mice exhibited very low levels of freezing which was significantly elevated during the POST-Shock period F_(1, 40)_ = 233.179, p = 0.000 [main effect (ME) of TIME: 3-way RM ANOVA] and increased with each successive shock F_(3, 120)_ = 105.876, p = 0.000 (ME of SHOCK: 3-way RM ANOVA; Figure 2C&G). ODOR did not have an effect on freezing during fear-conditioning (Figure 2B&F). Comparable levels of freezing were observed in hM4Di-Saline animals (Post-Shock 43.43 +/− 3.81%. Mice were then returned to the fear-conditioning chamber either 1d (RECENT) or 21d (REMOTE) later to assess fear memory recall. In animals that were tested for memory recall 1d after fear-conditioning (RECENT), inhibiting cell bodies in the dCA1 transfected with DREADDs did not have any behavioural consequences (Figure 2D) as there were no group differences or interactions in freezing levels [between-subject (BS) effects of ODOR & VIRUS: *ns*; 2-way ANOVA]. The Recall session lasted 5 minutes and all animals froze significantly less in the first minute compared to the rest of the session F_(1,95)_ = 5.127, p < 0.001 (ME of TIME: 3-way RM ANOVA; Figure 2E). Animals in the hM4Di-Saline group showed similar levels of freezing to the other groups (Total = 43.66 +/− 10.67%). In mice tested for recall at 21d (REMOTE), there was a decline in freezing in controls (NO ODOR, eYFP) and no difference between these animals and eYFP mice that received ODOR (Figure 2I). Surprisingly, both ODOR and NO ODOR mice that received DREADDs demonstrated modestly higher levels of freezing (Figure 2I) although this did not reach a level of statistical significance F_(1,21)_ = 4.133, p = 0.055 (ME of VIRUS: 2-way ANOVA). Similarly, when the session was divided into 1-minute bins, all mice froze significantly less in the first minute compared to the rest of the session F_(1,105)_ = 4.422, p = 0.002 (ME of TIME: 3-way RM ANOVA; Figure 2J) and DREADDs animals froze significantly more F_(1,105)_ = 14.742, p < 0.001 (ME of VIRUS: 3-way RM ANOVA; Figure 2J).

**Figure 2.**
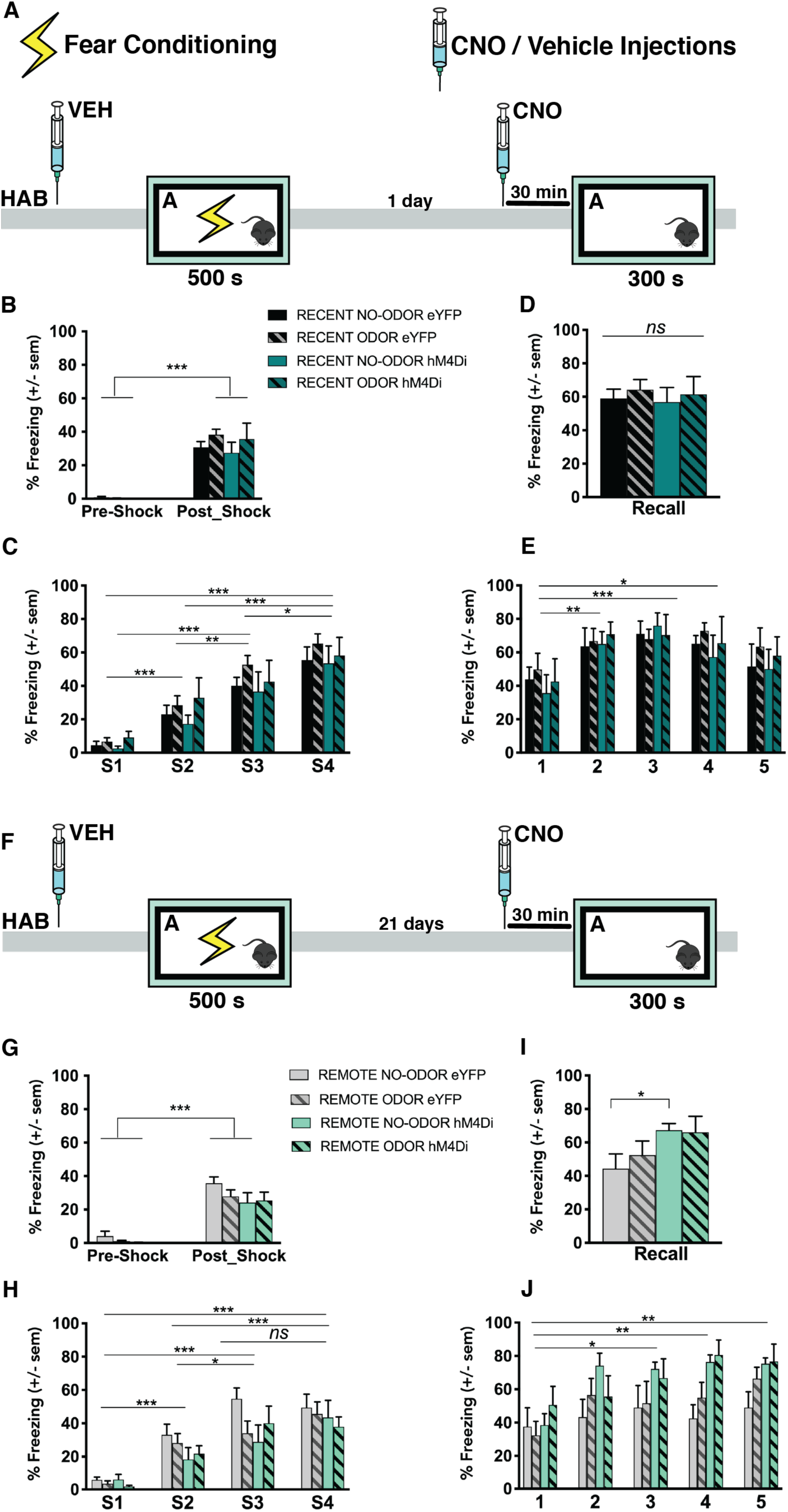
Inhibiting the CA1 with DREADDs did not Impair Fear Memory Recall. To assess whether odor modulates systems consolidation of a fear memory we employed a four-shock contextual fear conditioning (FC) protocol where animals were tested either in the presence of odor or no odor and tested for recall 1d (Recent) or 21d (Remote) later. **(A)** Timeline of our behavioral experiment for animals tested at 1d. **(B)** Animals were initially placed in context A without shock for 198s (Pre-Shock) where they exhibited very low levels of freezing which was significantly elevated during the Post-Shock period **(C)** and increased with each successive shock. **(D)** Mice were then returned to the FC context 1d later for a 5 min fear memory recall test. **(E)** Data shown across 1 min bins. Inhibiting cell bodies in the dCA1 with DREADDs did not have any behavioural consequences. **(F)** Timeline of our behavioral experiment for animals tested at 21d. **(G)** Animals were initially placed in context A without shock for 198s (Pre-Shock) where they exhibited very low levels of freezing which was significantly elevated during the Post-Shock period **(H)** and increased with each successive shock. **(I)** Mice were then returned to the FC context 21d later for a 5 min fear memory recall test. **(J)** Data shown across 1 min bins. Inhibiting cell bodies in the dCA1 with DREADDs caused an increase in freezing in both odor and no odor conditions. p< 0.05, ** = p<0.01, *** = p<0.001.

### Odor Modulates the Temporal Characteristics of Systems Consolidation

#### *c-Fos* Expression during Fear Memory Recall

Ninety minutes after the Recall session all animals were perfused so that *c-Fos* levels in the dCA1 and PL could be quantified. Given that inhibitory hM4Di DREADDs should theoretically silence neurons, and thus *c-Fos* expression, our predictions involving *c-Fos* expression are with respect to eYFP animals only. In line with SC theory, we hypothesized that REMOTE compared to RECENT fear memory recall would be less HPC-dependent evinced by lower *c-Fos* levels in the dCA1 at 21d compared to 1d and that the reverse profile would be observed in the PL. We also hypothesized that ODOR would bias this effect such that a fear memory associated with an odor would continue to be HPC-dependent even at the REMOTE time point.

#### Dorsal Hippocampus: dCA1

Figures 3E & 3I shows that in the dCA1, odor did not change the level of *c-Fos* expression at the RECENT time point as both ODOR and NO ODOR animals demonstrated similar *c-Fos* levels. In agreement with SC, we found that in eYFP animals, *c-Fos* levels declined at the REMOTE time point and as we predicted, specifically in the NO ODOR group and not in the ODOR group F_(1,32)_ = 4.173, p = 0.049 (ODOR × VIRUS interaction: 3-way ANOVA). Post-hoc analyses showed a significant difference between the NO ODOR eYFP groups at 1d and 21d t_(11)_ = 3.017, p = 0.012 and between eYFP ODOR and no ODOR groups at 21d t_(11)_ = 2.655, p = 0.022. Figures 3B & 3E shows that in mice conditioned with ODOR, immunoreactivity of *c-Fos* remained elevated even at the REMOTE time point F_(1,18)_ = 5.656, p = 0.029 (ODOR × VIRUS interaction: 2-way ANOVA) suggesting that odor can modulate the temporal characteristics of memory consolidation in that it can delay, or even potentially abolish the redistribution of fear memories from the HPC to the PFC.

**Figure 3.**
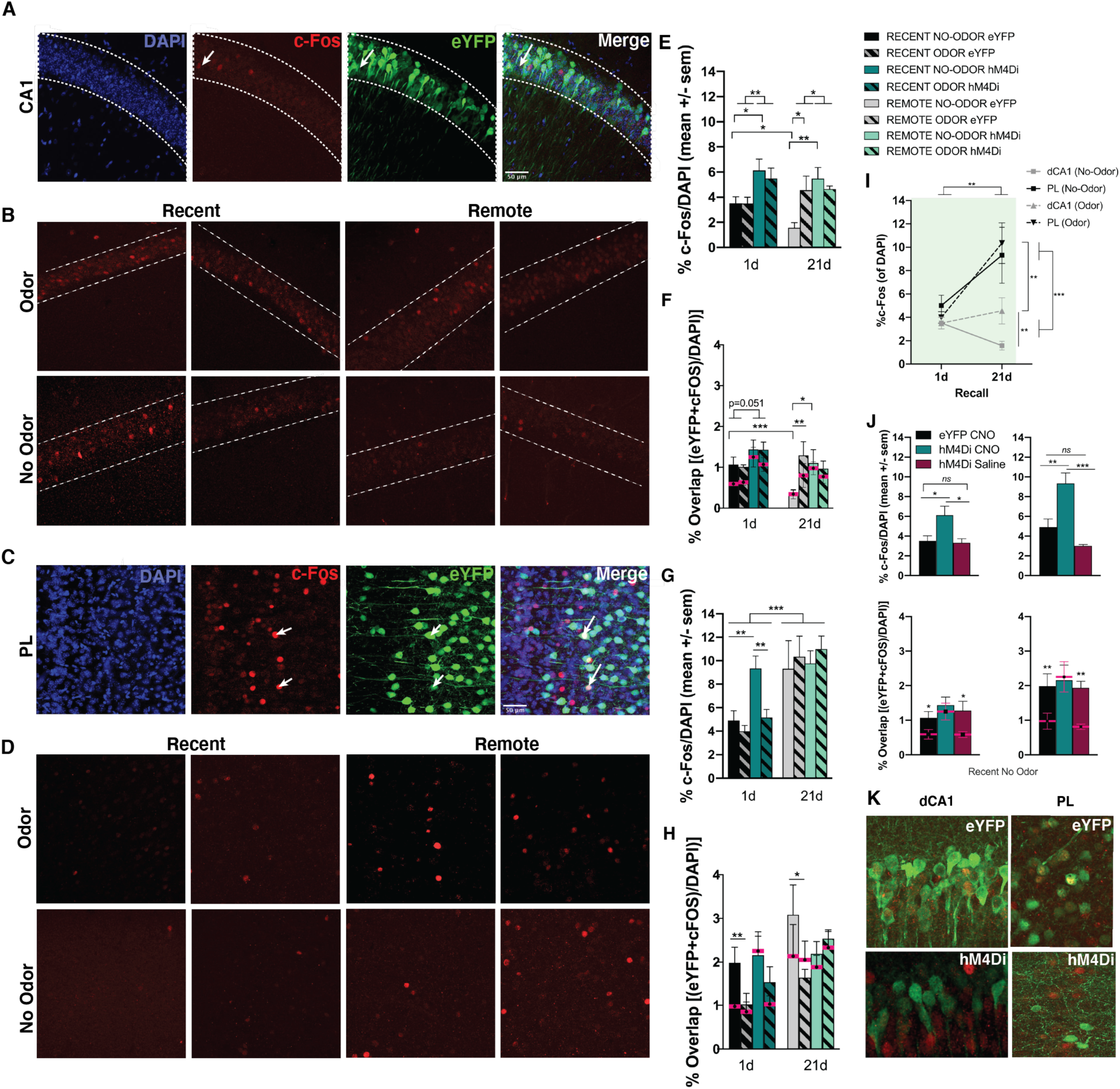
Odor Modulates the Temporal Characteristics of Systems Consolidation. For both recall sessions (1d & 21d), the total number of DAPI positive (+), eYFP+ (cells tagged during fear conditioning), *c-Fos*+ (cells active during fear memory recall), and eYFP+ and *c-Fos*+ (cells active in both behavioral epochs) neurons in the **(A)** dCA1 and **(C)** PL were quantified. **(B)** In eYFP animals, in dCA1 odor did not change *c-Fos* expression at 1d as predicted, *c-Fos* levels declined at 21 d in the no odor group but remained elevated in the odor group. **(D)** In eYFP animals, in the PL, *c-Fos* levels were significantly higher at 21d compared to 1d. Two images per condition are provided. **(E)** The % of *c-Fos* immunoreactive neurons was defined as a proportion of total DAPI labeled cells. In dCA1, there was a significant difference between no odor eYFP groups at 1d and 21d and between eYFP odor and no odor groups at 21d. Unexpectedly, *c-Fos* levels were higher in DREADDs animals compared to eYFP animals. **(F)** Overlaps were considered the proportion of DAPI labeled cells that were both c-Fos+ and eYFP+. Chance overlap was calculated as the % of eYFP+ neurons x the % of *c-Fos*+ neurons over the total number of DAPI neurons (pink line). Overlaps followed the same pattern as *c-Fos*. **(G)** In the PL, *c-Fos* levels were significantly elevated at 21d compared to 1d with the exception of the no odor group at 1d. As in dCA1, *c-Fos* was also higher in DREADDs animals compared to eYFP animals at 1d but not 21d. **(H)** Odor decreased overlaps to chance. **(I)** In eYFP animals, contextual fear memories become more reliant on the PL with time irrespective of odor. Conversely, they become less reliant on the dCA1 with time, if no odor is present. However, if odor is present during encoding and again upon retrieval, these memories continue to engage the dCA1. Inhibiting dCA1 with CNO using doxycycline-regulated, *c-Fos* promoter driven DREADDs introduced using an adeno-associated virus to tag fear engram cells produced an increase in *c-Fos* levels. **(J)** In the no odor groups at 1d, DREADDs + CNO mice showed greater numbers of *c-Fos+* cells, compared to DREADDs + Saline or eYFP + CNO mice in both the dcA1 (top left) and the PL (top right). These increases in *c-Fos* were predominantly seen in non-engram cells. When we view the proportion of double-labeled cells, they occur to a significantly higher degree that chance in the eYFP-CNO and hM4Di-Saline mice, but not for hM4Di-CNO animals where *c-Fos* is increased in non-engram cell populations in the dCA1 (bottom left) and the PL (bottom right). Pictured in **(K)** dCA1 (left) and the PL (right). p< 0.05, ** = p<0.01, *** = p<0.001.

At the RECENT time point there were no differences between the ODOR and NO ODOR groups F_(1,14)_ = 0.220, p = 0.646 (2-way ANOVA). However, unexpectedly *c-Fos* levels were higher in DREADDs animals compared to eYFP animals F_(1,32)_ =15.665, p < 0.001 (ME of VIRUS: 3-way ANOVA) and this was true at 1d F_(1,18)_ = 11.166, p = 0.005 (ME of VIRUS: 2-way ANOVA) and 21d F_(1,18)_ = 6.205, p = 0.023 (ME of VIRUS: 2-way ANOVA) (Figure 3E). This effect was even more pronounced in the NO ODOR condition at 1d q_(2)_ = 3.974, p = 0.014 and 21d q_(2)_ = 5.125, p = 0.002.

#### Prefrontal Cortex: PL

In Figures 3G & 3I, we found additional support for SC in the PL where *c-Fos* levels were significantly elevated at the REMOTE compared to RECENT time point F_(1,33)_ = 19.466, p < 0.001 (ME of RECALL: 3-way ANOVA). As in the dCA1, *c-Fos* levels in the PL were higher in DREADDs animals compared to eYFP animals at 1d but not at 21d F_(1,16)_ = 11.585, p = 0.004 (ME of VIRUS: 2-way ANOVA).

At the REMOTE time point, *c-Fos* levels were equally elevated for all groups with no effect of ODOR (Figure 3G & 3I). However, at the RECENT time point, *c-Fos* levels were elevated in DREADDs animals but only in the NO ODOR group F_(1,16)_ = 10.318, p = 0.005 (ME of ODOR: 2-way ANOVA, Figure 3G). Post hoc tests showed a significant difference between the NO ODOR DREADDs groups and the NO ODOR eYFP group q_(2)_ = 5.360, p = 0.002, and the ODOR DREADDs group q_(2)_ =4.718, p=0.004 at the RECENT time point. This suggests that inhibiting or perturbing the dCA1 with DREADDs during fear memory recall after 1d results in increased activity in the PL unless an ODOR is present. This further supports our hypothesis that ODOR can shift the temporal organization of fear memories to be less dependent on the PFC and more dependent on the HPC.

### Chemogenic Inhibition of Engram Cells Activates c-Fos in Neighbouring Non-Engram Cells

#### Fear Memory Encoding and Recall Overlaps

We next examined the overlap between labeled fear engram neurons (eYFP) and cells active during the Recall session (*c-Fos*). We hypothesized that in the dCA1, in NO ODOR animals, there would be higher overlap at the RECENT *vs.* REMOTE time point whereas in the ODOR animals we expected to see high overlap at both time points. We hypothesized that the opposite would be true in the PL with higher overlap at the REMOTE *vs.* RECENT time point and that ODOR would decrease this overlap. Again, our predictions involving overlap are with respect to eYFP animals only taking into consideration that we used inhibitory DREADDs to silence neurons. However, given that DREADDs in this experiment, increased *c-Fos* expression, we also looked at overlap in the DREADDs groups to assess whether we could glean insight to the increased levels of freezing seen at the REMOTE time point in these animals.

#### Dorsal Hippocampus: dCA1

In the dCA1, exhibited by a significant ODOR × RECALL interaction F_(1,19)_ = 5.486, p = 0.03 (2-way ANOVA), in accordance with our hypothesis, we found that within the NO ODOR eYFP condition there was a higher degree of overlap between the cells active during fear-conditioning and recall at the RECENT time point compared to the REMOTE time point t_(11)_ = 3.404, p = 0.006. This is in contrast to the ODOR eYFP condition where overlap was elevated at both time points (Figure 3F). A Mann-Whitney test indicated that overlap was greater in the ODOR group (Mdn = 1.057) compared to the NO ODOR group (Mdn = 0.262) at the REMOTE time point (U = 3, p = 0.008). Moreover, both ODOR and NO ODOR eYFP groups had higher than chance levels of overlap at the RECENT time point q_(2)_ = 4.03, p = 0.008 as well the ODOR eYFP group at the REMOTE time point q_(2)_ = 6.417, p < 0.001) (Figure 3F). For animals that received DREADDs, there were no differences in overlap across time points, and while the percentage of overlap was similar to eYFP animals, contrastingly, the percentage of overlap did not differ from chance as it did in the eYFP mice (Figure 3F) except for the ODOR group at the RECENT time point q_(2)_ = 3.861, p = 0.01. In the NO OODR animals, overlap here was greater in the DREADDs group compared to the eYFP group at the REMOTE time point q_(2)_ = 3.186, p = 0.037. This pattern suggests that *c-Fos* levels were increased in neighbouring non-engram cells (Figure 3K left).

#### Prefrontal Cortex: PL

In the PL, a three-way ANOVA revealed a significant main effect of RECALL F_(1,33)_ = 6.808, p = 0.014 and a significant main effect of ODOR F_(1,33)_ = 6.787, p = 0.014 as well as a significant ODOR × VIRUS interaction F_(1,33)_ = 4.382, p = 0.044. Specifically within the eYFP animals using a two-way ANOVA, there was a significant main effect of RECALL F_(1,19)_ = 4.697, p = 0.043 and a significant main effect of ODOR F_(1,19)_ = 9.874, p = 0.005 but no interaction signaling that ODOR decreased overlap to chance (only eYFP NO ODOR groups differed from chance: RECENT – q_(2)_ = 5.211, p < 0.001; REMOTE – q_(2)_ = 4.499, p = 0.003, all other groups *ns*) at both time points (Figure 3H). This suggests that odor-associated memories are not processed in the PL or at least suggests that this is the case when recall occurs in the presence of the same odor that was present at the time of encoding. Post-hoc analyses revealed that this is certainly true at the RECENT time point q_(2)_ = 3.15, p = 0.033 specifically for eYFP animals q_(2)_ = 4.414, p = 0.007. However, due to high levels of variability in the NO ODOR group this effect was not observed at the REMOTE time point q_(2)_ = 2.08, p = 0.151 (Figure 3H). A two-way ANOVA across ODOR and VIRUS at the REMOTE time point revealed a significant difference between the ODOR and NO ODOR groups in the eYFP condition q_(2)_ = 3.937, p = 0.013. It remains unknown whether overlap would still be low if encoding had been carried out in the presence of an odor as was the case in the current experiment, but the recall test had been conducted in the absence of odor.

Given that *c-Fos* levels were higher in the dCA1 in DREADDs animals compared to eYFP animals t_(14)_ = −4.112, p = 0.001 (Figure 3B) but that overlap was higher in the eYFP animals compared to the DREADDs animals at the RECENT time point F_(1, 14)_ = 4.539, p = 0.051, 2-way ANOVA) and the REMOTE time point F_(1,18)_ = 4.563, p = 0.047 (ODOR × VIRUS interaction, 2-way ANOVA), this suggests that in the dCA1, DREADDs increased *c-Fos* and not necessarily in fear memory neuronal ensembles (Figure 3K) and may be primarily increasing activity in neighboring neurons in a compensatory manner. We ran an additional control experiment without odor at the RECENT time point. We included a group (hM4Di Saline) that was injected with the hM4Di virus in the dCA1 (as well as the eYFP virus in the PL), however, these animals received saline instead of CNO. We compared *c-Fos* levels in these animals to mice that were given CNO as well as injected with the hM4Di virus (hM4Di CNO) or the eYFP virus (eYFP CNO), in the dCA1 (as well as the eYFP virus in the PL). Saline injections resulted in comparable expression of *c-Fos* in the dCA1 to eYFP animals (F_(2,13)_ = 5.753, p = 0.019, one-way ANOVA) (Figure 3J, top left). When we look at overlaps compared to chance we see that the increases *c-Fos* occur within dCA1 engram cells for the eYFP-CNO (t_(3)_ = 3.910, p = 0.011) and hM4Di-Saline animals (t_(3)_ = −3.535, p = 0.039), but not for hM4Di-CNO animals where *c-Fos* is increased in non-engram cell populations (Figure 3J, bottom left) suggesting that chemogenetically silencing the neurons involved in fear memory encoding during recall 1d later, can result in increased *c-Fos* activity in non-engram hippocampal cells.

Interestingly, *c-Fos* levels were also increased in the PL in DREADDs animals, with the exception of the ODOR group at the RECENT time point (Figure 3G). However, overlap was only elevated above chance for the NO ODOR groups at both RECENT and REMOTE time points suggesting that even in the PL, DREADDs seem to be increasing *c-Fos* in a compensatory manner following inhibition of dCA1 engram cells, and not in PL engram cells (Figure 3K). Moreover, when we looked at *c-Fos* levels in the PL in the control animals that had received saline instead of CNO, we found that they were also significantly lower than hM4Di-CNO animals demonstrating that inhibiting fear memory engram cells can cause an increase in activity in the PL (F_(2,13)_ = 4.84, p = 0.031, one-way ANOVA) (Figure 3J, top right). When we look at overlaps compared to chance we see that the increases *c-Fos* occur within PL engram cells for the eYFP-CNO (t_(5)_ = −6.263, p = 0.002) and hM4Di-Saline animals (t_(3)_ = −7.903, p = 0.004), but not for hM4Di-CNO animals where *c-Fos* is increased in non-engram cell populations (Figure 3J, bottom right).

### Freezing was Positively Correlated *with c-Fos Expression and Overlap in the dCA1 and Negatively Correlated with c-Fos Expression in the PL in Controls*

We ran bivariate correlational analyses between freezing levels during the Recall session and *c-Fos* expression as well as Overlap in the dCA1 and in the PL. In NO ODOR eYFP animals we found that freezing was correlated with *c-Fos* (r = 0.729, n = 11, p=0.011, 2-tailed) and Overlap (r = 0.612, n = 11, p = 0.046, 2-tailed) in the dCA1 (Figures 4A-B) and negatively correlated with *c-Fos* in the PL (r = −0.647, n = 9, p=0.032, 2-tailed) (Figures 4C-D). When this was parsed across time, we saw no significant correlations. These animals showed a decrease in freezing across time and also a decrease in *c-Fos* levels in the dCA1 as well as an increase in *c-Fos* in the PL. In NO ODOR DREADDs animals, freezing during Recall did not correlate with *c-Fos* or Overlap in either brain region. In ODOR animals, we saw no significant correlations in eYFP or DREADDs animals and when we looked specifically at the groups that demonstrated increased freezing at the REMOTE time point, their freezing levels were not correlated with *c-Fos* or Overlap in either the dCA1 or the PL. This suggests that there is a dissociation between these measures and freezing as a behavioral output, specifically in animals that received DREADDs and therefore, the increases in *c-Fos* observed in this condition are not likely related to the increased freezing observed at the REMOTE time point. However, it is possible that there was a ceiling effect on freezing reached here and these cells which are now c-Fos+ due to DREADD inhibition are conceivably contributing to freezing.

**Figure 4.**
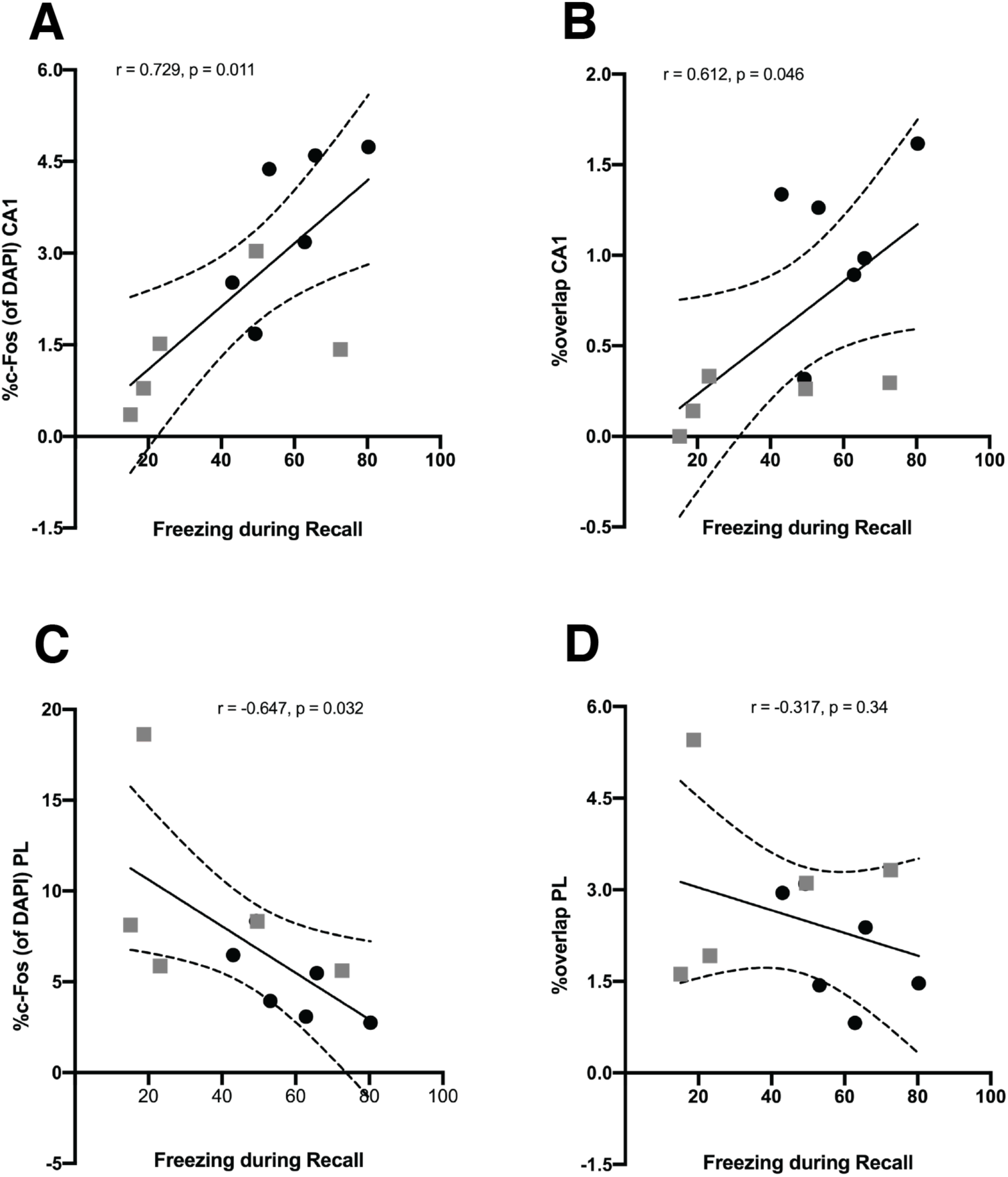
Freezing was Positively Correlated with c-Fos Expression and Overlap in the dCA1 and Negatively Correlated with c-Fos Expression in the PL in Controls. We ran bivariate correlational analyses between freezing levels during the recall sessions and *c-Fos* expression as well as overlap in the dCA1 and in the PL. In no odor eYFP animals we found that freezing was correlated with **(A)** *c-Fos* and **(B)** overlap in the dCA1 and negatively correlated with **(C)** *c-Fos* in the PL. **(D)** There was no correlation between freezing and overlap in the PL. p< 0.05, ** = p<0.01, *** = p<0.001.

## DISCUSSION

As the French author Marcel Proust described long ago (Proust, 1927), referenced in his work *A la Recherche du Temps Perdu* which translates to *In Search of Lost Time* - “Yet a single scent already breathed long ago, may once again both be in the present and the past, be real without being present”. Sensory experiences like olfaction and gustation have the ability to produce the evocation of vivid autobiographical memories of the distant past (Toffolo et al., 2012; Zucco et al., 2012; Daniels and Vermetten, 2016; Glachet and El Haj, 2019). This spontaneous process which we have all experienced at one time or another is often referred to as the Proust Phenomenon. Few studies have tried to unravel this phenomenon systematically. For most, this reverie can be a pleasant trip through time but for individuals with PTSD, odors can serve as triggers for extremely aversive experiences (Toffolo et al., 2012). The vivid nature of these experiences can make them all the more painful.

In this study we sought to determine whether odor as a contextual cue, associated with a fear-related memory, could shift the temporal dynamics of the consolidation of that memory at the systems level. To assess this, we fear conditioned animals using a four-shock protocol either with or without almond odor present within the training context. We observed increased freezing during training with each successive shock and that odor did not affect the overall learning curves. The lack of group differences during conditioning also suggests that there were no intrinsic effects of the hM4Di virus in the absence of CNO. We then replicated previous work showing that with the passage of time, fear memories are processed to a lesser extent within the HPC and rely on neocortical storage (Teixeira et al., 2006; Lesburgueres et al., 2011; Silva et al., 2018; DeNardo et al., 2019). When fear memory recall following conditioning was tested after 1 d, we observed a more pronounced recruitment of cellular activity compared to the PL. In contrast, when tested after 21 d, we detected increased activity in the PL compared to the dCA1. These data support SC in that episodic-like memory retrieval recruits cortical structures such as the PL at remote compared to recent time points, whereas the HPC is important for initial memory storage (Frankland and Bontempi, 2005).

Fear conditioning has been widely used as a model for PTSD in rodent studies (Rasmusson and Charney, 1997; Morrison and Ressler, 2014; Bali and Jaggi, 2015), and fear memories are encoded as engrams across multiple brain regions (Josselyn et al., 2015) and activation of these engrams has been shown to be both necessary and sufficient to drive fear responses. However, while the foundation for cortical traces are laid down early during encoding (Lesburgueres et al., 2011; Wang et al., 2012; Cardenas et al., 2019; Corches et al., 2019; Jacques et al., 2019) there is evidence to suggest that the PFC neurons involved in fear memory retrieval are not the same neurons active during encoding (Giannotti et al., 2019; DeNardo et al., 2019). Within the PFC, we specifically looked at the PL for several reasons. In humans, the dorsal anterior cingulate cortex (ACC) has been shown to be hyperactive in response to fearful stimuli in individuals with PTSD (Milad et al., 2007) and comparatively in rodents the anatomic equivalent of the ACC is the PL (Choi et al., 2010; Laubach et al., 2018). In rodent studies, the PL has been implicated in the expression and renewal of fear responses (Corcoran and Quirk, 2007; Sierra-Mercado et al., 2011; Orsini and Maren, 2012; Courtin et al., 2014; Fenton et al., 2014; Corches et al., 2019) and increased activity in this region is associated with extinction deficits (Burgos-Robles et al., 2009) as well as increased input to the basolateral amygdala (BLA) (Likhtik et al., 2005; Maren et al., 2013), an output which promotes freezing behavior and fear (Vidal-Gonzalez et al., 2006).

We tested whether presenting an odor during fear conditioning, and then again during retrieval, would alter the dynamics of consolidation in a way that engaged the HPC for a longer period of time (e.g. high intensity training has been shown to have the opposite effect, speeding up the decay of HPC dependency promoting memory generalization; Pedraza et al., 2017). We found that odor influenced this temporal relationship biasing the memory system away from the PFC and towards the HPC, suggesting that retrieval of highly contextual experiences does require the HPC even at remote time points. When recall was tested after 1 d, *c-Fos* levels were equally elevated in dCA1 in both odor and no-odor groups. However, when we looked at recall at 21 d, these levels declined in the no-odor group but not in the odor animals. Overlaps were also higher and statistically above chance, in the dCA1 at the recent time point for both odor and no odor groups and at the remote time point when odor was present, suggesting that odor can mediate how fear memories are organized.

The present study did not include a group that was exposed to odor during encoding but not during retrieval, opening up the possibility for future experiments to test whether odor as a function of memory possesses the ability to delay translocation of the fear memory trace to the PFC, or if odor presented at the time of retrieval re-contextualizes the memory activating the HPC as a result. This data would provide insight to the nuances within the existing debate regarding theories of consolidation: is the HPC always engaged when processing a highly contextual memory (they decontextualize slower) as purported by MTT? Or are these memories equally subject to context generalization, schematization, and decontextualization but given the ideal cue to evoke recall can become recontextualized thereby stimulating the HPC? Future experiments may also include a group that tests this as well measurements linking *c-Fos* expression to context generalization (Wiltgen and Silva, 2007). To further support the modulatory role of odor on systems consolidation, we showed that when odor was present, overlaps within the PL decreased to chance levels at both time points, suggesting that odor-associated memories recruit different neurons from encoding to retrieval. It would also be interesting to measure whether overlaps would remain at chance levels if odor had been presented during encoding but not retrieval, as there is a possibility that the PL processes the odor as a new cue reverting the system back to encoding, despite the observed bias for odor to be processed in the HPC.

Olfaction is a unique sensory modality in that it is the only sense that bypasses the thalamus sending projections directly to the forebrain (Shepherd, 2005). Olfactory input is a highly salient sensory cue in both rodents and humans, and guides many different types of behavior from maternal bonding and reproduction, to social hierarchy, foraging, and the detection of pathogens (Doty, 1986; Kesslak et al., 1988; Edwards et al., 1990; Fleming et al., 1999; Li et al., 2007; Lübke and Pause, 2014; Takahashi et al., 2018). Qualitatively, this may explain why odor as a contextual cue possesses seemingly time-travelling characteristics, (Tulving, 2002) where one whiff of your mother’s favorite perfume can bring you back to your childhood in an instant. Odor is tightly interconnected with memory. The dentate gyrus (DG) of the HPC is important for keeping highly similar memories and experiences separate, a function called pattern separation (Sahay et al., 2011). It has been shown that the ventral DG plays an important role in olfactory learning and memory, and specifically in odor discrimination (Weeden et al., 2014) which may help elucidate how odor-related memories are so resistant to interference. Additionally, there is evidence to suggest that individuals with memory disorders such as Alzheimer’s disease experience deficits in processing olfactory memory (Kesslak et al., 1988), potentially due to its reliance on the HPC.

Finally, we assessed whether inhibiting the tagged cellular ensembles in the dCA1 using DREADDs would decrease freezing. Given the upregulation of the HPC at the recent time point compared to the remote time point we hypothesized that inhibiting this region would affect memory recall only at 1 d and not at 21 d. In eYFP controls, we demonstrated that odor promotes hippocampal involvement during remote memory recall, therefore, we also hypothesized that inhibiting the dCA1 would impair memory at the remote time point if the memory was associated with an odor. Behaviorally, we saw no effect of inhibiting the dCA1 with CNO on recall at 1 d and unexpectedly there was an increase in freezing at 21 d. The incubation of fear hypothesis originally proposed by Hans Eysenck (Eysenck, 1968) states that repeated exposure to an unreinforced conditioned stimulus (i.e., the training context) following acquisition of a classical aversive conditioned response (i.e., freezing) provided that the conditioned stimulus (i.e., the shock) is a strong stimulus, will serve to enhance the conditioned response over time (Richards and Martin, 1990). In previous rodent studies, which have examined fear memory recall across recent and remote time points, fear responses related to the training context tended to either stay stable (Wiltgen and Silva, 2007; Poulos et al., 2016) or strengthen over time (Houston et al., 1999; Balogh et al., 2002; Frankland et al., 2004; Poulos et al., 2016) supporting Eysenck’s hypothesis. However, there have also been human studies where this effect was not observed (Richards and Martin, 1990), and in the current study we saw a decline in freezing over time. While we did not test fear generalization, it is possible that this decline in the conditioned response was accompanied by an increase in generalization (Wiltgen and Silva, 2007). If our animals had not shown this decrease, we speculate that the behavioral differences between eYFP and DREADDs animals at the remote time point would not have been observed. The majority of studies reporting incubation of fear looked at recall between 28-36 days later (Balogh et al., 2002; Frankland et al., 2004; Wiltgen and Silva, 2007; Poulos et al., 2016; Germer et al., 2019) with one study (Siegmund and Wotjak, 2007) claiming maximal fear expression occurred at 28 days, whereas we tested our mice at 21 days which may not be long enough for an effect of fear incubation to emerge. Interestingly, Houston et al., (1999) found that fear incubation is less pronounced in older rats (27 months). Together, the conclusions from our behavioural data should be tempered nonetheless, chemogenic inhibition of dCA1 engram cells using DREADDs did not decrease freezing at either time point.

Unexpectedly, we also saw high levels of *c-Fos* expression in all groups that received DREADDs in the dCA1. Theoretically, these neurons should effectively be silenced and show very little, if any *c-Fos* expression. DREADDs are G-protein coupled receptors (GPCRs) that have been genetically engineered to allow minimal constitutive activity when they are introduced by viral-mediated gene transfer directly into the brain, thus behavior in the animal is not affected by their presence (Pei et al., 2008). DREADDs do not have endogenous ligands as they are activated only by the synthetic ligand, CNO, an inert compound that does not induce any other activity in the rodent brain aside from binding to these designer GPCRs (Pei et al., 2008). The inhibitory DREADDs construct that we used contained the gene hM4Di (Gi) which originated from the human muscarinic receptor where induced point mutations allow inefficacy of the endogenous ligand acetylcholine, and efficacy of CNO, where activation of hM4Di receptors can inhibit neurons in a reversible manner (Armbruster et al., 2007). Confirmed using extracellular *in vivo* recordings measured 30 min later, CNO administered at 5mg/kg was able to inhibit neuronal firing rates in *c-Fos*-tTA transgenic mice injected with an AAV virus carrying the hM4Di gene under the control of the hSyn1 promoter (Ryan et al., 2015). Following this time frame, as DREADDs are reversible, firing rates rebounded back to steady state. Several other electrophysiology studies have produced similar findings (Armbruster et al., 2007; Ferguson et al., 2011; Parnaudeau et al., 2013; Sano et al., 2014). For the most part, other studies which have established the inhibitory nature of this gene have tested whether CNO sufficiently hyperpolarizes neurons in slice preparations with a CNO bath-application onto cultured hippocampal cells *in vitro.* The degree to which CNO can hyperpolarize the cell has been shown to be dose dependent (Zhu et al., 2014). It is possible that the dose we chose (3mg/kg) was too low to observe behavioral effects, and potentially our dose was not sufficient to cross the threshold of decreased firing to silence neurons but resulted in “quieting” neurons rather than shutting them off completely. However, many other studies that have reported observable effects employed lower doses (1mg/kg) (Andero et al., 2014; Mahler et al., 2014; Sweeney and Yang, 2015; Parfitt et al., 2017; Chiang et al., 2018).

We have explored several possible explanations for our DREADD results. The most parsimonious is that CNO itself produced a cellular perturbation that did not fully manifest into a behavioral readout. Thus, although we failed to see a behavioral effect, we nevertheless observed alterations in *c-Fos* expression. It is also possible that our virus targeted interneurons and therefore rather than silencing excitatory pyramidal cells in the dCA1, we disinhibited inhibitory interneurons causing an increase in activity. Theoretically, this fits with our behavioral results since further increases in activity in the dCA1 would not have contributed to further freezing, as a ceiling effect was potentially reached. In contrast, in *c-Fos*-tTA transgenic mice (Reijmers et al., 2007) transduced with the DOX-regulated AAV9-TRE-ChR2-eYFP virus, engram labeling was restricted to excitatory neurons, as no overlap was detected somatically between ChR2 positive cells and inhibitory gamma-aminobutyric acid (GABA) positive cells (Liu et al., 2012). To date, no assessment of the kinetics of CNO based neural inactivation has been conducted in wildtype animals that have been injected with the *c-Fos*-tTA-TRE-hM4Di-eYFP virus, although in a recent paper our group too observed similar results of *c-Fos* expression in the BLA (Chen et al., 2019).

One of the striking differences between the DREADDs system we used and what has been used in the majority of other studies is that our system is a tet-off system where hM4Di is only expressed in a subset of cells transduced (∼20%). Targeted cells specifically consist of the fear engram cells active during tagging / encoding and not all of the neurons in that particular brain region. In a recent paper, using *c-Fos-*tTA transgenic mice injected with a DOX-regulated TRE-ArchT-GFP virus to optogenetically silence cocaine-place engram neurons in the dCA1, light delivery led to inhibition in the targeted neurons accompanied by the simultaneous emergence of non-targeted, alternative neurons in the vicinity. They found that their manipulation caused a disengagement from initially recruited cells (during encoding) and the activation of an alternative spatial map (Trouche et al., 2016). Although this study was conducted using optogenetics, it is possible that silencing engram cells in the dCA1 resulted in the recruitment of adjacent non-engram cells. This would explain how *c-Fos* were elevated, but overlaps were not statistically above chance, following CNO injections in hM4Di mice. Furthermore, promoter-dependent differential hM4Di virus expression has been observed (Lopez et al., 2016).

It is not surprising that we would see these compensatory, off-target effects following acute perturbation of the steady state internal dynamics of the brain possibly interfering with the excitatory-inhibitory balance in the dCA1 (Otchy et al., 2015). This is colloquially known as the “whack a mole” effect, once you hit a mole another one pops up. In this case silencing a neuron is like hitting a mole. We also saw an increase in *c-Fos* in the PL following inhibition in the dCA1. This was observed in all groups except the group tested after 1 d that was exposed to odor. Again, given the complex excitatory-inhibitory feedback networks within the HPC and between the HPC and the PFC, inhibition or perturbation in the dCA1 with DREADDs would not necessarily result in a decrease in firing in the PL (Schmidt et al., 2019). Schmidt et al., (2019) found that some neurons increased, and some neurons decreased their firing rate following CNO injection in mice transfected with the hM4Di gene in the PFC under the CAMKIIa promoter. Interestingly, in the current study, perturbing the dCA1 at the recent time point redirected processing during memory retrieval to the PL in the no-odor group but not the odor group. This further supports our hypotheses and suggests that to some degree odor-associated fear memories are less malleable in the sense that they require the HPC for processing. We believe that, together, this presents a putative new target for PTSD research. Current therapeutic interventions often involve exposure therapy, and reactivating a previously consolidated, odor-associated fear memory in the presence of the associated odor may bias the memory system allocating the trace to the HPC for processing. Theoretically, this may help focus intervention strategies to a particular locus or node in the brain that can be combined with other treatments. Additionally, future studies may involve intersectional analysis involving this theory and the use of optogenetics.

Previous research has demonstrated a connection between the expression of excitatory DREADDs and *c-Fos* (Garner et al., 2012; Roy et al., 2019). To assess whether there was a link between increases in *c-Fos* expression observed following inhibition of the dCA1 with DREADDs and our behavioral data where we saw an increase in freezing in animals that received CNO at the remote time point, we ran a correlational analysis. Several studies have shown a positive correlation between conditioned freezing and *c-Fos* expression in the HPC (Radulovic et al., 1998; Strekalova et al., 2003; Knapska and Maren, 2009), however, some studies have shown a dissociation (Singewald, 2007; Plendl and Wotjak, 2010; Mastrodonato et al., 2018). We only found significant correlations in our eYFP no odor group where conditioned freezing was positively correlated with *c-Fos* levels and the percentage of overlap in the dCA1 and negatively correlated with *c-Fos* expression in the PL. Since we found no correlations between these measures in the DREADDs groups, the increase in freezing that we saw in the DREADDs mice at the remote time point cannot be attributed to the increased *c-Fos* expression seen in these mice in the dCA1 and in the PL following CNO administration. Moreover, the time interval between CNO injection and perfusion was 120 minutes, and it remains possible that by this time *c-Fos* levels started to rebound following neuronal inhibition and that we would have seen lower *c-Fos* levels and significant correlations if we had sacrificed the animals slightly earlier.

In summary, our histological results provide insight into the molecular dynamics related to activity-dependent, DOX-regulated inhibitory DREADDs, while also demonstrating plasticity in the process of memory consolidation at the systems level whereby even remote memories are amenable to modulation by contextual cues such as odor. A better understanding of how memories are layered, comprised of contextual information at the level of the engram localized to specific brain regions (Vetere et al., 2019) may provide therapeutic insight to the convergence of emotional processing and PTSD symptomology (Daniels and Vermetten, 2016).

## MATERIALS & METHODS

### Animals

51 wild type male c57BL/6 mice (2-3 months of age; Charles River Labs, Wilmington, MA) weighing 18-25g at the time of arrival were housed in groups of 2-5 mice per cage. Mice were kept on a regular light cycle 12:12 hour light/dark in a temperature and humidity-controlled colony room. Cages were changed once a week and contained cardboard huts and nesting material for enrichment. Upon arrival in the facility, mice were placed on a 40 mg/kg doxycycline (DOX) diet (Bio-Serv, product F4159, Lot 226766) and left undisturbed for a minimum of 3 days prior to surgery with access to food and water *ad libitum*. They were then handled for two consecutive days for 2 minutes each day. The following day they underwent surgery. All subjects were treated in accordance with protocol 17-008 approved by the Institutional Animal Care and Use Committee at Boston University. The behavioural results reported are based on data from a total of 51 experimental animals. Due to a loss of tissue histological data reported is based on 44 animals for CA1 and 45 animals for PL.

### Stereotaxic Surgery and Virus Microinjections

Aseptic surgeries were carried out with mice mounted into a stereotaxic frame (Kopf Instruments, Tujunga, CA) with skull flat. They were anesthetized with 4% isoflurane and 70% oxygen for induction with isoflurane at 2% to maintain anesthesia during surgery. A small amount of 2% lidocaine (Clipper Distributing Company, St. Joseph, MO) was placed on the skull as a topical analgesic and a small hole was drilled above each injection site. Animals received four microinjections, bilateral injections into the dorsal CA1 (dCA1) (Fig 1A-B) and the prelimbic cortex (PL) (Fig 1A, C) via a 10ul gas-tight Hamilton syringe attached to a micro-infusion pump (UMP3, World Precision Instruments, Florida, USA) which occurred at a rate of 100 nl min^−1^. Coordinates are given relative to Bregma (in mm). All mice were injected bilaterally in the PL and the dCA1. In the PL, mice were injected with a viral cocktail of AAV9-cFos-tTa (titre: 1E+13GC/mL) and TRE-eYFP (titre: 3E+13GC/mL) in a volume of 400 nl/side at AP: +1.9, ML: +/− 0.3, DV: −1.9. The dCA1 was targeted at AP: −2.0, ML: +/− 1.5, DV: −1.2, with mice receiving a cocktail of AAV9-cFos-tTa and either TRE-hM4Di-eYFP (titre: 2.5E+13GC/mL) (DREADDs) or TRE-eYFP (eYFP) in a volume of 550 nl/side. Injectors were left in place for 1 minute following each injection to avoid liquid backflow. Mice were then sutured, received 0.2ml physiological sterile saline (0.9%, s.c.) and 0.1 mL of a 0.03 mg/ml buprenorphine solution (i.p.) at the end of the surgery, and were placed on a heating pad and given hydrogel. Mice were given an additional injection buprenorphine (0.1 mL, 0.03 mg/ml, i.p.) the next day. Except for cage changes, having their tails were marked with a skin marker every 24-48 hours, and being weighed, mice were left undisturbed for a ten-day period following surgery to allow for recovery and virus expression.

### Behavioral Testing: Fear Conditioning and Recall

Mice were habituated to 2%DMSO injections in sterile saline (0.9%, i.p) 3 days prior to fear conditioning. DOX diet was replaced with standard rodent chow (*ad libitum*) 42 hours prior to behavioral tagging. Half of the animals were fear conditioned in the presence of almond extract-soaked gauze in a plastic container with holes (ODOR) in the conditioning chambers (Colbourn, Whitehall, PA) while the other half were just given the tubes without the odor (NO ODOR). The fear conditioning session lasted 500 seconds. Mice received four shocks during this session: 1^st^ shock at 198s, 2^nd^ at 280s, 3^rd^ at 360s, and 4^th^ at 440s (Costanzi et al., 2014; Redondo et al., 2014) (2s duration, 1.5 mA intensity). At the end of the session, mice were placed back on DOX (*ad libitum*) and placed in a holding tank until all cage mates had been fear-conditioned and then all mice were placed back with cage mates in a clean home cage. Mice were returned to the original fear-conditioning context for a Recall session either 1 day (RECENT) or 21 days later (REMOTE) with the same odor conditions they received during conditioning. Thirty minutes prior to the recall session, all mice were injected with clozapine-N-oxide (CNO) and 90 minutes after the recall session mice were euthanized. We also included a control group that was tested at the RECENT time point with NO ODOR, however these mice were given saline injections (0.9%, i.p.) instead of CNO (n=4). Each conditioning chamber had a camera mounted on the roof for video recording. Video was fed into a computer running Freeze Frame / View software (Coulbourn Instruments) where freezing was defined as a bout of immobility lasting 1.25 seconds or longer.

### Drugs: Use of Clozapine-N-Oxide

Half of the animals were injected with an inhibitory DREADDs virus (hM4Di) fused to an eYFP reporter while the other half were given just eYFP. To inhibit neuronal firing (Pei et al., 2008; Dong et al., 2010; Ferguson et al., 2011; Zhu and Roth, 2015; Roth, 2016) in cells transfected with DREADDs (Mahler et al., 2014; Zhu and Roth, 2015), all mice (except for the saline control group) were injected with CNO, obtained from NIH: NIDA Drug Supply Program (NIMH C-929; batch ID: 14073-1) and prepared in a concentration of 0.6 mg/ml in sterile saline (0.9%) and 2% DMSO. Thirty minutes prior to the Recall session this was administered i.p. at a dose of 3 mg/kg.

### Immunohistochemistry

Mice were overdosed with isoflurane and perfused transcardially with (4° C) phosphate-buffered saline (PBS) followed by 4% paraformaldehyde (PFA) in PBS. Brains were extracted and stored overnight in PFA at 4°C and transferred to PBS solution the following day. Brains were sliced into 50μm coronal sections with a vibratome (Leica, VT100S) and collected in cold PBS. Sections were blocked for 2 hrs at room temperature in 1x phosphate-buffered saline + 2% Triton (PBS-T) and 5% normal goat serum (NGS) on a shaker. Sections were transferred to well plates containing primary antibodies made in PBS-T (1:1000 rabbit anti-*c-Fos* [SySy]; 1:5000 chicken anti-GFP [Invitrogen]) and incubated on a shaker at 4°C for 48 hrs. Sections were then washed in PBS-T for 10 min (x3), followed by 2 hr incubation with secondary antibody (1:200 Alexa 555 anti-rabbit [Invitrogen]; 1:200 Alexa 488 anti-chicken [Invitrogen] made in PBS-T). Following three additional 10 min washes in PBS-T, sections were mounted onto micro slides (VWR International, LCC). Nuclei were counterstained with DAPI added to Vectashield HardSet Mounting Medium (Vector Laboratories, Inc), slides were then cover-slipped and put in the fridge overnight. The following day the edges were sealed with clear nail-polish and the slides were stored in a slide box in the fridge until imaging.

### Fluorescent Confocal Image Acquisition & Analysis

Images were collected from coronal sections a fluorescent confocal microscope (Zeiss LSM 800 with airyscan) at 20X magnification. For each animal, in the PL, 2-3 z-stacks (step size 0.94um) were taken per hemisphere from 2-3 different slices yielding 4-6 total z-stacks per animal. In the dCA1, z-stacks (step size 0.94um) were obtained from both hemispheres from 3-4 different slices. Per slice two stacks were taken in the left hemisphere, one in the dorsomedial CA1 and one in the dorsolateral CA1. The values for these were summed. Two images were also taken in the right hemisphere, and these also summed. Data from each hemisphere was then pooled and the mean for the total 6-8 z-stacks were computed. The total number of DAPI positive (+) neurons were counted using Image J / Fiji (https://imagej.nih.gov/ij/). In addition, the number of eYFP+ (cells tagged during fear conditioning), *c-Fos*+ (cells active at the time of fear memory recall), and eYFP+ and *c-Fos*+ (cells active in both behavioral epochs) neurons in the dCA1 and PL were quantified to measure the number of active cells during defined behavioral tasks. Percentage of immunoreactive neurons was defined as a proportion of total DAPI labeled cells. Chance overlap was calculated as the percentage of eYFP+ neurons multiplied by the percentage of *c-Fos*+ neurons over the total number of DAPI neurons.

### Statistical Analysis

Calculated statistics are presented as means ±SEM. To analyze differences, we used three-way analyses of variance (ANOVA) [RECALL (between-subject factor - two levels: RECENT & REMOTE time points) × ODOR (between-subject factor – two levels: ODOR & NO ODOR) × VIRUS (between-subject factor – two levels: eYFP & DREADDS)]. When appropriate, follow-up comparisons (two-way ANOVAs, independent T-tests) and post-hoc analyses (Tukey’s HSD, or Mann-Whitney) were conducted. When Fear Conditioning and Recall sessions were divided into bins, these data were analyzed using Repeated Measures (RM) three-way ANOVAs [ODOR (between-subject factor - two levels: ODOR & NO ODOR) × VIRUS (between-subject factor – two levels: eYFP & DREADDS) × TIME (within-subject factor – two levels: PRE & POST Shock / or five levels: MINUTES 1-5) or × SHOCK (within-subject factor – four levels: SHOCK 1-4). Comparisons between the percentage of overlap against chance were also conducted using three-way RM ANOVAs [ODOR (between-subject factor - two levels: ODOR & NO ODOR) × VIRUS (between-subject factor – two levels: eYFP & DREADDS) × CHANCE (within-subject factor – two levels: OVERLAP & CHANCE). Bivariate correlational analyses were performed using Pearson’s Correlation Coefficient. All statistical tests assumed an alpha level of 0.05. For all figures, * = p< 0.05, ** = p<0.01, *** = p<0.001.

## ACKNOWLEDGEMENTS

We would like to thank Susumu Tonegawa and his lab for providing the activity-dependent virus cocktail. We would like to thank the NIH for providing us with CNO. We would also like to thank the members of the Ramirez lab for helpful comments and suggestions. This work was supported by NIH Early Independence Award (DP5 OD023106-01), a Young Investigator Grant from the Brain and Behavior Research Foundation, a Ludwig Family Foundation Grant, and the McKnight Foundation Memory and Cognitive Disorders Award.

## AUTHOR CONTRIBUTIONS

All authors contributed to the design of the experiments and data collection. S.L.G. and A.H.F. conducted all behavioral experiments, and S.L.G., A.H.F., O.M., and H.L. contributed to quantification of the histological data. S.L.G., A.H.F. and S.R. wrote the paper. All authors edited and commented upon the manuscript.

